# Effect of geographic isolation on the nasal virome of indigenous children

**DOI:** 10.1101/616342

**Authors:** Eda Altan, Juan Carlos Dib, Andres Rojas Gulloso, Duamaco Escribano Juandigua, Xutao Deng, Roberta Bruhn, Kristen Hildebrand, Pamela Freiden, Janie Yamamoto, Stacey Schultz-Cherry, Eric Delwart

## Abstract

The influence of living in small remote villages on the diversity of viruses in the nasal mucosa was investigated in three Colombian villages with increasing levels of geographic isolation. Viral metagenomics was used to characterize viral nucleic acids on nasal swabs of 63 apparently healthy young children. Sequences from human virus members of the families *Anelloviridae, Papillomaviridae, Picornaviridae, Herpesviridae, Polyomaviridae, Adenoviridae, and Paramyxoviridae* were detected in a decreasing fraction of children. The diversity of human viruses was not reduced in the most isolated indigenous Kogi villages. Multiple viral transmission clusters were also identified as closely related variants of rhinoviruses A or B in 2 to 4 children from each of villages. The number of papillomavirus detected was greater in the village most exposed to outside contacts while conversely more anellovirus infections were detected in the more isolated indigenous villages. Genomes of viruses not known to infect humans, including in the family *Parvoviridae* (genus densoviruses)*, Partitiviridae, Dicistroviridae, and Iflaviridae and* circular Rep expressing ssDNA genomes (CRESS-DNA) were also detected in nasal swabs likely reflecting environmental contamination from insect, fungal, and unknown sources. Despite the high level of geographic and cultural isolation, the diversity of human viruses in the nasal passages of children was not reduced in indigenous villages indicating ongoing exposure to globally circulating viruses.

**Importance:** Extreme geographic and cultural isolation can still be found in some indigenous South American villages. Such isolation may be expected to limit the introduction of globally circulating viruses. Very small population size may also result in rapid local viral extinction due to lack of sufficient sero-negative subjects to maintain transmission chains of rapidly cleared viruses. We compared the viruses in the nasal passage of young children in three villages with increasing level of isolation. We find that isolation did not reduce the diversity of viral infections in the most isolated villages. Ongoing viral transmission of rhinoviruses could also be detected within all villages. We conclude that despite their geographic isolation remote villages are continuously exposed to globally circulating respiratory viruses.

## Introduction

The impact of geographic isolation in shaping the respiratory virome remains largely unknown. In the pre-agricultural era, people typically lived widely dispersed in small nomadic groups, a lifestyle which may have minimized the spread and maintenance of infectious diseases that did not establish long lasting or chronic infections. Small populations now settled in hard to explore regions may still be relatively isolated from repeated exposures to highly prevalent viruses circulating in larger, more connected, communities. Inhabitants of such highly isolated villages may have therefore lost viruses dependent on large population size of young, seronegative, susceptible hosts found in larger populations [1].

Coincident with the arrival of Europeans, native Amerindian populations underwent strong population bottlenecks possibly due to imported airborne epidemics such as small pox, measles, and more recently influenza viruses to which they had no prior exposure [2, 3]. To determine whether reduced rate of outside contact coincides with a reduction or even an absence of detectable human viruses, we analyzed and compared the nasal virome of children in two highly isolated Amerindian Kogi villages in a tropical forest of Northern Colombia and of one largely Hispanic village alongside a coastal highway. In order to detect all human viruses, viral metagenomics was applied to nasal swabs collected from children two to nine years old.

## Results

### Sample collection and village location

Nasal swabs were collected from 63 children (53.9% female) with a mean age of 5 years (Table 1). The children lived in three Northern Columbia villages that differed in degree of outside contacts. Samples used for comparison were from age, sex and race matched children (Table 1). The first village Calabazo (GPS 11.28448, −74.00195) is located along a major road (highway 90) running alongside the National Natural Park of Tayrona and is frequently visited by tourists. Calabazo has a 2005 census population size of 499 and the main language is Spanish. Seywiaka (GPS 11.2174, −73.5794) is an isolated village with a population size of 250-300 accessible only by foot (1.5 hours walk from nearest road) inhabited by Kogi people speaking their indigenous language (Fig 1A). An even more isolated Kogi village Umandita (GPS 11.09698,−73.64781) with an estimated population of 350-400 inhabitants is accessible after a 9-10 hours walk from Seywiaka (Fig 1B).

**Table 1.** Age and gender of children analyzed.

|  | 2 to 5 years old | 6 to 9 years old | Girl | Boy |
| --- | --- | --- | --- | --- |
| Calabazo | 11 | 10 | 8 | 13 |
| Seywiaka | 12 | 9 | 9 | 12 |
| Umandita | 13 | 8 | 17 | 4 |

**Figure 1A.**
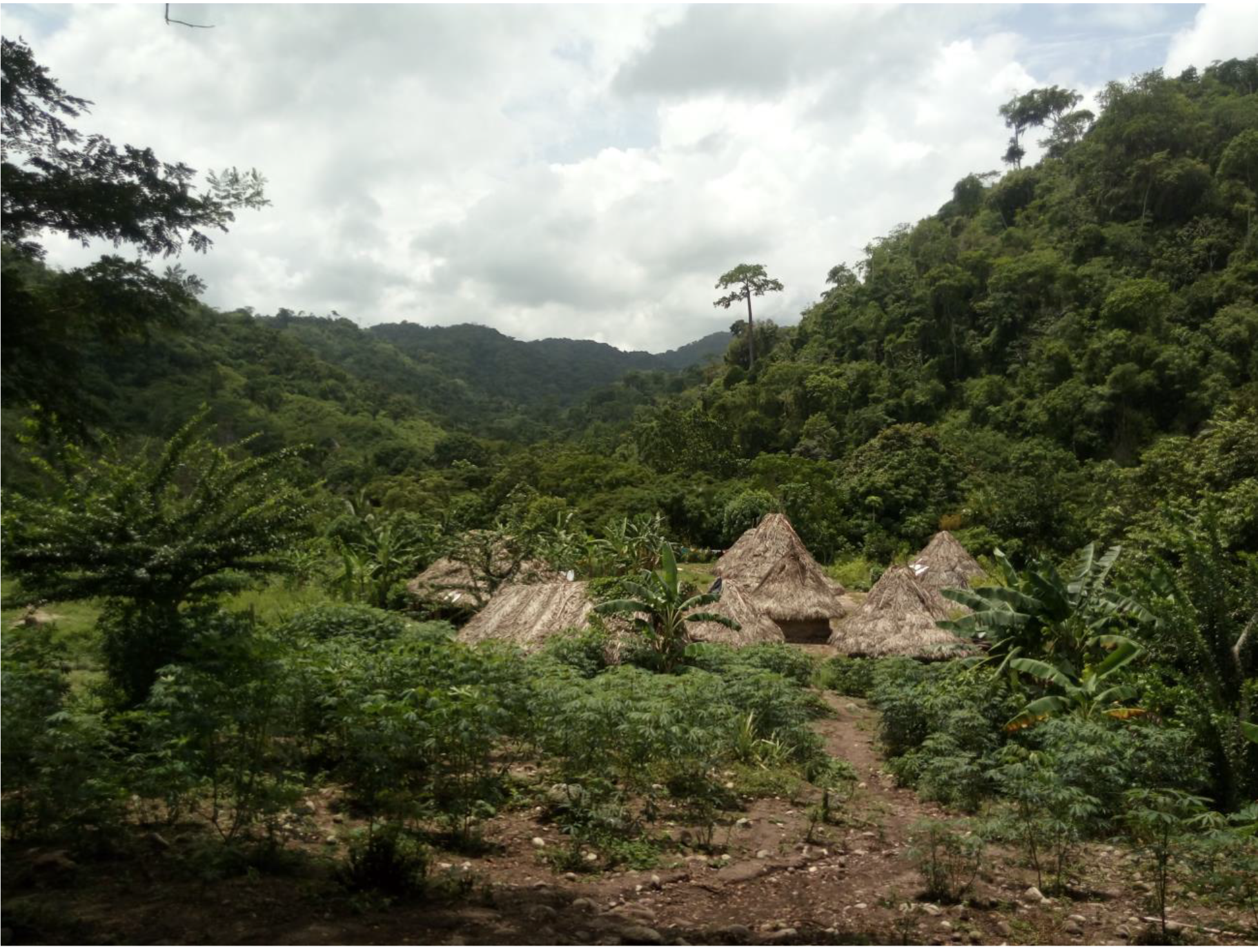
View of Seywiaka village.

**Figure 1B.**
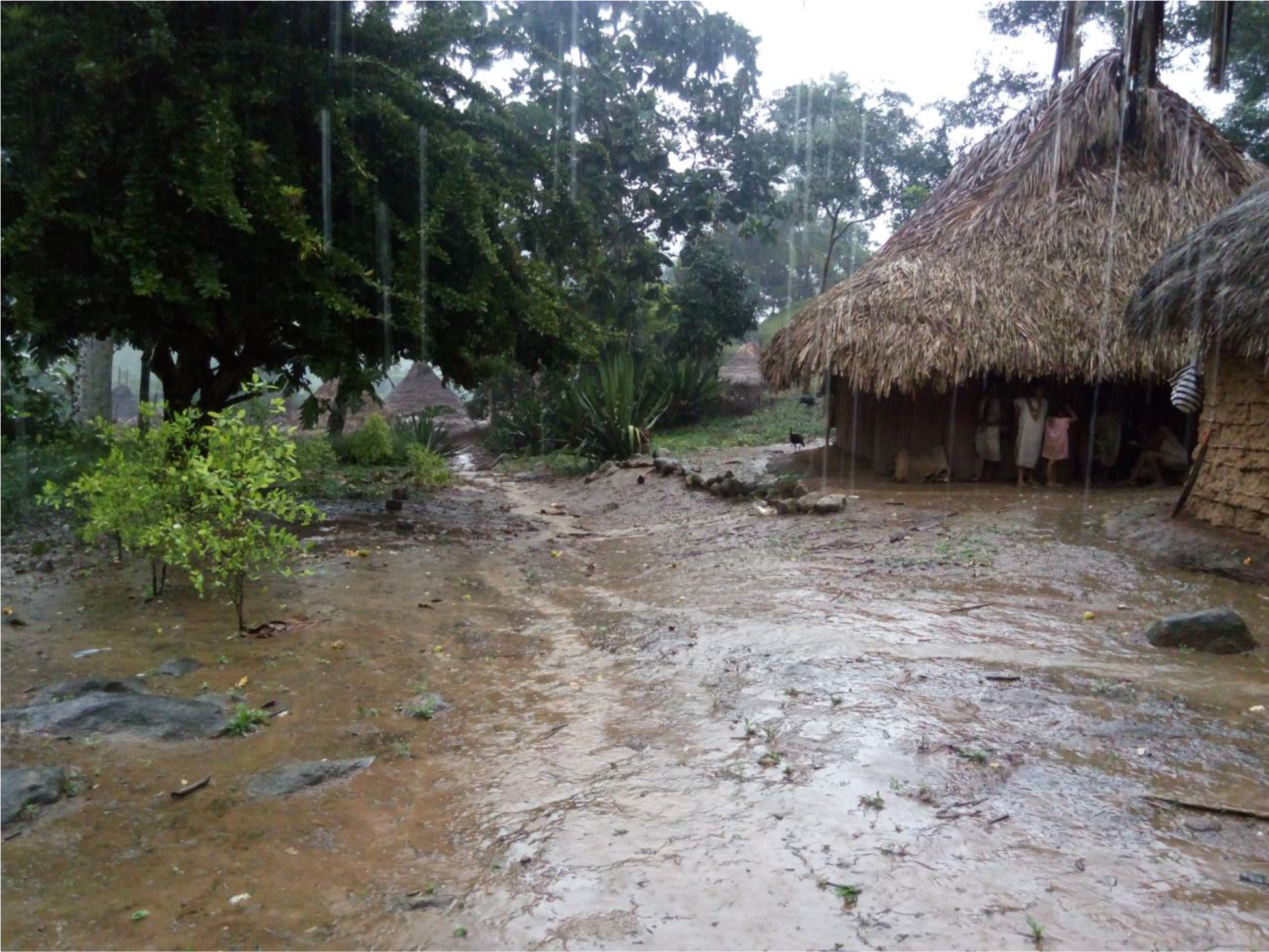
View of Umandita village.

### Nasal mucosa virome

Following viral metagenomics enrichment of viral particles-associated nucleic acids in nasal swabs, random RNA and DNA amplification, and deep sequencing of 63 individual nasal swab supernatants a total number of 63 million reads were generated for an average number of reads of approximately one million per sample. The raw sequence data for each pool is available at NCBI’s Short Reads Archive under GenBank accession number PRJNA530270. We found 92% of samples (58/63) to be positive for at least one human virus. Human virus belonging to 7 viral families were detected and are listed in decreasing prevalence of detection (*Anelloviridae, Papillomaviridae, Picornaviridae, Herpesviridae, Polyomaviridae, Pneumoviridae, and Adenoviridae*).

Anelloviridae family members reads were the most commonly detected viral sequences and were found in 49/63 children or 77.7%. 0.16% (n=100,957 sequence reads) of 63 million total reads could be mapped to the *Anelloviridae* family with BLASTx E scores <10^−10^. The second most commonly detected human virus reads belonged to the *Papillomaviridae* family, which were detected in 44.4% (28/63 children) with 0.087% of total reads (n=55,248). Next, with a prevalence of 23.8% (15/63 children) were reads from the *Picornaviradae* family. Of these, 0.094% reads (n=59,819) encoded picornavirus reads from the species Rhinovirus A (10/63 children, 15.8%), Rhinovirus B (3/63 children, 4.7%), Rhinovirus C (1/63 children, 1.58%) and Enterovirus B (1/63 children, 1.58%). *Herpesviridae* family members were next in prevalence being detected in 7/63 children (11.1%) including human betaherpesvirus 5 (CMV or HHV5)(6/63 children, 9.52%), human herpesvirus 6 (Roseolovirus or HHV6) (3/63 children, 4.76%), and human betaherpesvirus 7 (Kaposi Sarcoma virus or HHV7) (1/63 children, 1.58%). In the *Polyomaviridae* family, human polyomavirus 5 (Merkel Cell carcinoma virus or HPyV5 [4]) (1/63 children, 1.58%), human polyomavirus 10 [5,6], (1/63 children, 1.58%), human polyomavirus 11[7] (1/63 children, 1.58%) were detected. Adenovirus C reads were detected in 2/63 children (3.17%). Respiratory syncytial virus (RSV), belonging to the *Paramyxoviridae* family, was found in 2/63 children (3.17%), This was the only viral family detected exclusively in the most exposed Calabazo village. The fraction of total reads from each sample encoding proteins with high-level similarity (E scores <10^−10^) to human viruses are shown in (Fig. 2) with the exception of the papillomaviruses and anelloviruses that are analyzed below.

**Figure 2.**
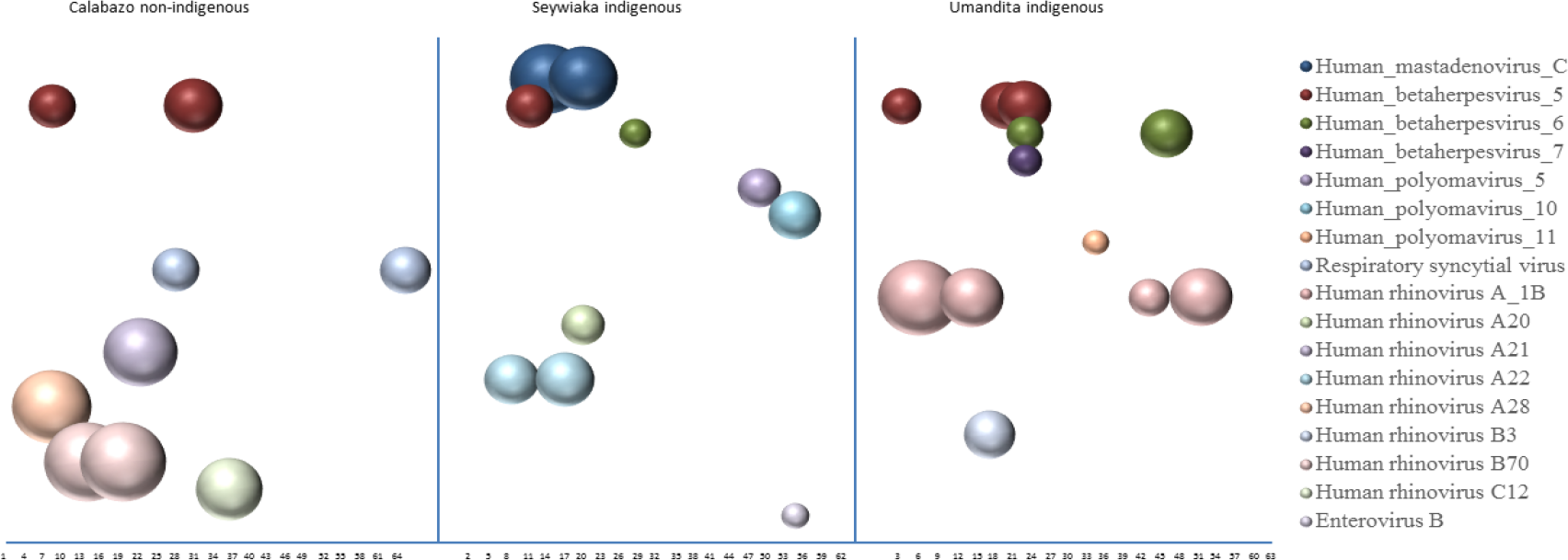
Distribution and level of viral reads to human viruses in the three villages.

### Family *Picornaviridae*

Fourteen children showed the presence of picornavirus sequences, 13 from rhinoviruses A, B, or C species and 1 from enterovirus B species. Rhinovirus (RV) reads generated contigs ranging in size from 481 to 7,089 nt (GenBank accession no MK501733 - MK501745). In total, 8 contigs included complete 5UTR-VP4-VP2, 2 contigs complete 5UTR-VP4, and 1 contigs a complete VP4-VP2 sequences that were used for phylogenetic analysis (Fig 3). Four rhinovirus contigs from Umandita region showed closest nucleotide identity (90 to 92%) to genotype 1B of rhinovirus A (RV-A-1B). Contigs from 3 of these children overlapped over almost the entire genome (6.6Kb without gaps) and showed a nucleotide identity of 99.8-99.9%. These three rhinovirus A-1B contigs clustered tightly together reflecting a recent common origin and an ongoing transmission cluster in the most isolated village, Umandita. Two children from Calabazo were infected with rhinovirus B70 that had 99.9% nucleotide identity indicating another transmission cluster occurring at the time of sampling. Two children from Seywiaka were shedding rhinovirus A22 but did not generate enough sequence reads to be included in phylogenetic analysis. Reads from these two children did overlap by 154 bases showing a single mismatch indicating another possible transmission cluster. Rhinovirus transmission clusters (genotypes A1B in Umandita, B70 in Calabazo, and A22 in Seywiaka) were therefore detected in each village. The enterovirus B reads from a Seywiaka child showed closest amino acid identity (93%) to Echovirus E15 (GenBank AY302541).

**Figure 3.**
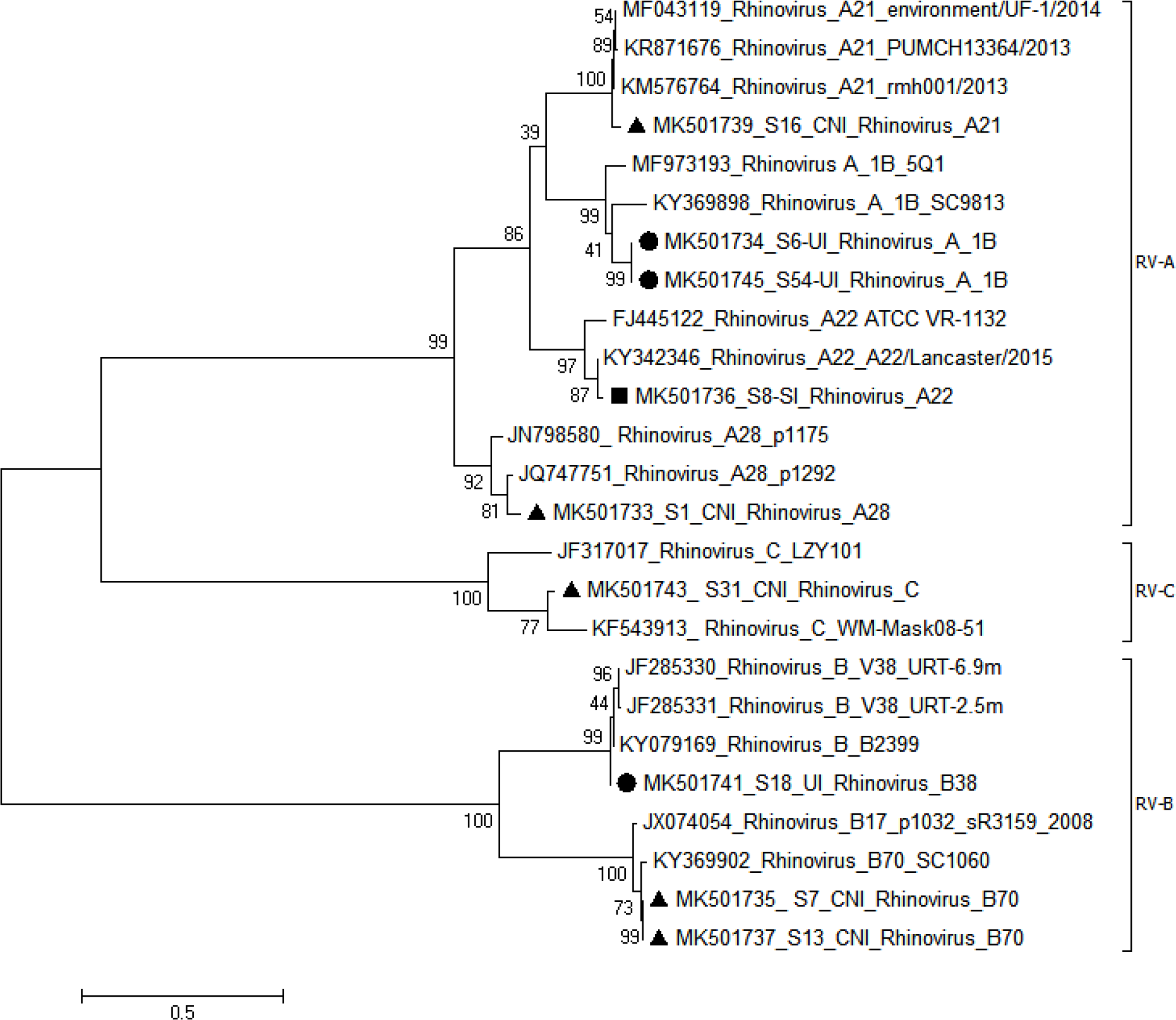
Phylogenetic analysis of VP4-VP2 region of rhinoviruses. ● Calabazo non-indigenous (CNI), ▲Seywiaka indigenous (SNI), ▄Umandita indigenous(UI)

### Polyomaviruses

Polyomavirus sequences were also found in the isolated villages of Seywiaka (n=2) and Umandita (n=1). Two Seywiaka villagers were shedding human polyomavirus 5 (Merkel cell polyomavirus), or human polyomavirus 10 (MW polyomavirus), and one child from Umandita was shedding human polyomavirus 11 (STL polyomavirus)(Fig 2).

### Herpesviruses

Sequences of human CMV, roseolovirus, and Kaposi sarcoma virus were identified. CMV sequences were found in six children (2, 1 and 3 children from Calabazo, Seywiaka and Umandita respectively). Three children shed Roseolovirus (2 and 1 from Seywiaka and Umandita respectively). One Kaposi sarcoma virus infection was detected in a child from Umandita (Fig 2). All contigs showed 98-100% nucleotide identities to genomes in GenBank.

### Adenovirus, pneumovirus, and parvovirus

Sequences from human_mastadenoviruses C species (HAdV-C) in the *Adenoviridae* family ranging in size from 250 to 1831 nt, were identified in two children from Seywiaka village (Fig 2). Six different regions (E3, E4, E1a, and L3) showed overlap between children with nucleotide identity of 83 to 97% likely reflecting two independent infections

Two respiratory syncytial virus generating contigs of size 363 nt and 888 nt were generated from two children in Calabazo both showing 99% nucleotide identity with respiratory syncytial virus strain A (GB accession number MG793382) (Fig 2). Contigs from these two children overlapped in the G gene (350 bp) showing a nucleotide identity of 99.1%. The close sequence identity of these two RSV strain may also reflect an ongoing transmission cluster within that village.

Unexpectedly, two reads of canine bocavirus were also detected in one swab sample from Calabazo village showing 92 and 97% aa identity to canine bocavirus NS1 gene region (GB accession number MG025952).

### Anelloviruses

Reads matching *Anelloviridae* family viruses were found in 77.7% (49/63) of children. Prevalence of anellovirus detection was 42% (10/21), 90% (19/21) and 95.2% (20/21) in children from Calabazo, Seywiaka and Umandita respectively. The overall fraction of children infected with different anellovirus genera were 34% with alphatorquevirus, 44.4% with betatorquevirus, 28.5% with gammatorquevirus and 65% with unclassified anelloviruses.

### Papillomaviruses

Altogether we detected 29 papillomaviruses consisting of 17 genotypes in 13 Calabazo children; 10 papillomaviruses consisting of 9 genotypes in 9 Seywiaka children; and 6 papillomaviruses consisting of 6 genotypes in 6 Umandita children (Figure 4B).

**Figure 4A and B.**
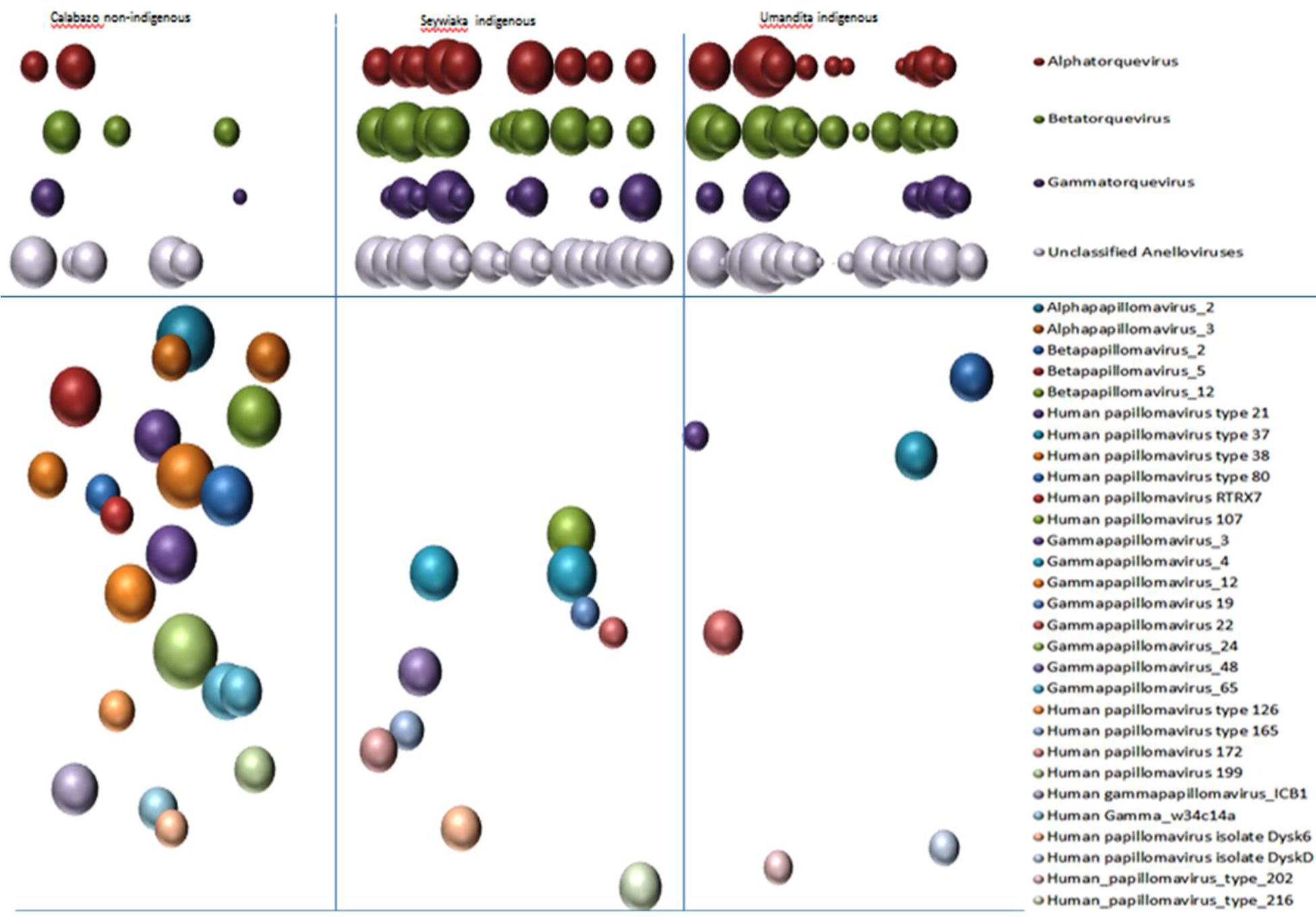
Distribution and level of anellovirus and papillomavirus reads in all three villages.

37 partial papillomavirus genome contigs ranging in size from 261 nucleotides (nt) to 7,392 nt were generated, 14 of which included a partial L1 gene region. Phylogenetic analysis of these L1 sequences was generated (Fig 4). All papillomavirus contigs showed 97-100% aa identities to papillomavirus proteins in GenBank. Papillomaviruses (HPV12) in two children from Calabazo village (S13-CNI, S49-CNI), were closely linked (Fig 5) showing 99% nucleotide identity.

**Figure 5.**
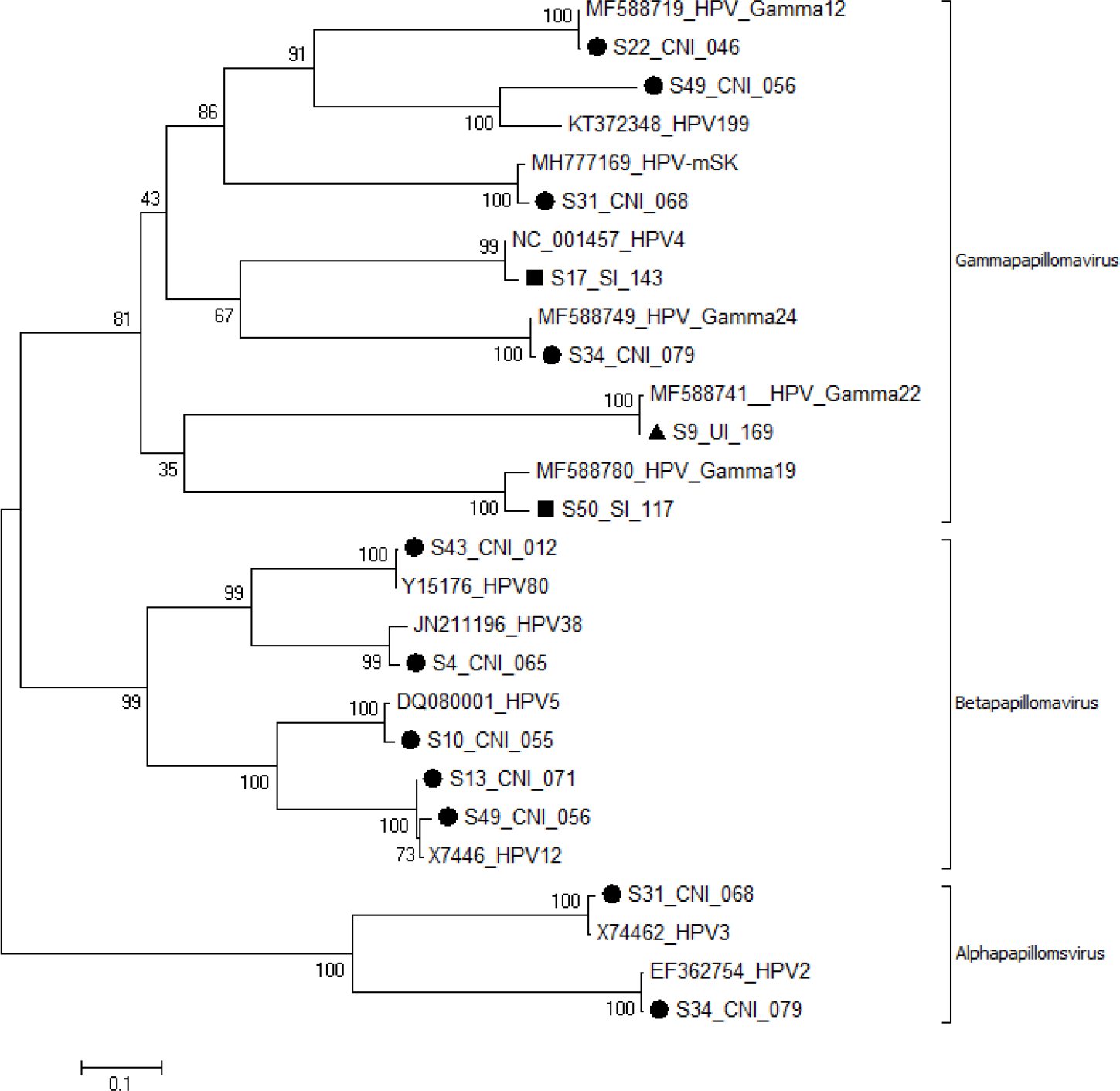
Phylogenetic analysis of major capsid (L1) protein of papillomaviruses. ● Calabazo non-indigenous (CNI), ▲Seywiaka indigenous (SNI), ▄Umandita indigenous(UI)

### Virome comparison between villages

We next compared the distribution of the two viral families that yielded the most reads, anelloviruses and papillomaviruses, among the 3 villages (Fig 4). The numbers of anellovirus infections were significantly different among the villages (p=0.0001). Inspection of the anellovirus distribution indicated that fewer infections were detected in the most exposed Hispanic village of Calabazo.

An analysis of papillomavirus reads distribution also showed that the number of papillomavirus infections were significantly different among the villages (p=0.043). As opposed to anelloviruses a greater number of papillomavirus infections were detected in the two most isolated villages relative to Calabazo (Fig 4).

### Viral families of non-vertebrate or unknown host tropism

Sequences from viral families not known to infect human (or vertebrates), likely representing air-borne mucosal surfaces contamination, were detected in 44/63 (69.8%) of children. Members of the following viral clades, ranked from highest to lowest prevalence, were detected (*Parvoviridae*-densoviruses, *Partitiviridae, Dicistroviridae, circular Rep encoding single stranded DNA viruses-CRESSS-DNA, and Iflaviridae)* were found in 49.2%, 38.09%, 30.1%, 23.8%, 4.7% of the swab samples respectively (Fig 6).

**Figure 6.**
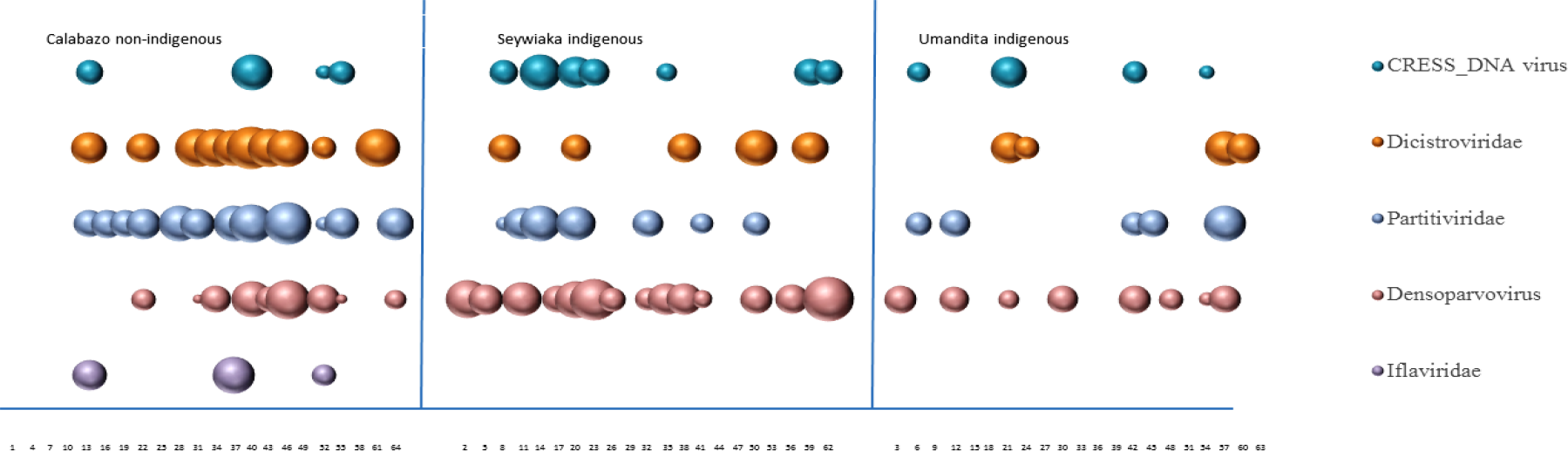
Distribution of viral sequences from viral groups not known to infect vertebrates or of unknown tropism.

Densoparvoviruses, dicistroviruses, and iflaviruses are known to infect invertebrates and partitiviruses to infect fungi and plants. Some *CRESS-DNA* viruses can infect fungi, plants, or mammals, but the tropism of most CRESS-DNA genomes largely identified through metagenomics of environmental samples (including those detected here) remain unknown.

## Discussion

Viral metagenomics of human respiratory secretions have analyzed different sample types mainly from clinical cases while only a few studies have studied samples from healthy subjects. Early studies of nasopharyngeal aspirates from lower respiratory tract infections in China [8] and Sweden [9] revealed numerous viruses with members of the families *Paramyxoviridae* (respiratory syncytia virus, metapneumovirus, parainfluenza virus), *Picornaviridae* (rhinovirus), *Orthomyxoviridae* (influenza) pre-dominating. A viral metagenomics analysis of nasopharyngeal swabs, aspirates, and sputums from patients with acute respiratory infections could confirm the presence of diverse viruses expected from PCR results [10]. Pediatric nasopharyngeal swabs viral metagenomics results also showed high concordance with the results of a commercial respiratory virus panel as well as reveal the presence of other, non-tested for, viruses [11]. Nasopharyngeal/oropharyngeal swabs from children with pneumonia of unknown etiology and asymptomatic controls, when analyzed by viral metagenomics could identify numerous viruses some of which could be associated with respiratory symptoms [12]. Nasopharyngeal swabs from children with community-acquired pneumonia but negative for common respiratory viruses revealed non-tested for viruses and human parainfluenza 3 virus with a large deletion that may have precluded specific PCR amplification [13] highlighting an advantage of non-specific amplification and deep sequencing. Lung transplant recipients with respiratory symptoms showed human rhinovirus infections recalcitrant to PCR detection as well as frequent HHV7 and anellovirus infections [14]. Diverse viruses could be detected in about a third of immune-compromised children with pulmonary disease including co-infections missed by prior clinical tests together with untested for viruses [15]. Metagenomics analyses therefore shows great promises as a supplement or even replacement for more specific viral genome detection assays although sensitivity issues remain [16–19].

A more limited number of metagenomics studies have analyzed the respiratory track virome of healthy children. Double stranded DNA from the anterior nares of healthy individuals in the human microbiome project showed that beta and gamma papillomaviruses were the most common viruses detected, followed by roseolovirus (HHV6) [20]. A PCR study of sinonasal mucosa from sinus surgery patients for 16 common respiratory viruses indicated HHV6 was the most the frequently detected virus [21]. A study of nasopharyngeal swabs from healthy 18 months old children showed the presence of human rhinoviruses, adenoviruses, bocaviruses, and parainfluenza virus [22]. Nasopharyngeal swabs from healthy children showed anelloviruses, HHV6, and HHV7 to be the most common infections [12].

Here we characterize the nasal mucosal viromes of 63 age and gender matched children from three villages and show that the geographical and cultural isolation of the two indigenous villages did not eliminate or even reduce the diversity of their human viruses. Three different herpesviruses (HHV5-7) and three different polyomaviruses (human polyomavirus 5, 10, 11) were detected in the two isolated villages while only one herpesvirus (HHV5) and no polyomaviruses were found in the village with frequent outside contact (Fig 2). Respiratory syncytia virus was the only virus found exclusively in the most exposed village. Four different rhinovirus and one enterovirus B genotypes were found in the two isolated villages while four rhinovirus genotypes were detected in the more exposed village. There was no overlap in the picornavirus genotypes in the different villages. In 0-5 years old children, the rates of asymptomatic rhinovirus detection have been reported from 12.5 to 33% [23–26]. In under 3 years old asymptomatic children, a rhinovirus detection rate of 33% did not significantly differ from that found in matched hospitalized children [25]. Here, we found an average 20.6% rate of rhinovirus detection in healthy 2-9 years old children, ranging from 23% in Calabazo and Umandita villages to 14% in Seywiaka.

Outbreaks, as reflected by the detection of closely related variants of the same rhinovirus genotypes, were detected in both the most isolated (four cases of rhinovirus A1B in Umandita) and the least isolated (two cases of rhinovirus B70 in Calabazo) villages. Two rhinovirus A22 infections in isolated Seywiaka were also very closely related and likely also epidemiologically linked. Because rhinovirus infections are of short duration it seems likely that each of these clusters resulted from recent introductions within these communities.

The origin of sequence reads from viral clades not known to infect humans, namely from the *Parvoviridae* genus densoparvovirus, *Partitiviridae, Dicistroviridae, Iflaviridae* and CRESS_DNA viruses remains unknown but deposition onto nasal mucosa surfaces from environmental sources such as the ambient air remains a likely possibility. Possible source for such viruses include plants and fungi for the partitiviruses, and insects for the dicistroviruses, iflaviviruses, and densoparvoviruses. The origins of CRESS-DNA viral genomes are unknown. The detection of a very few reads of canine bocavirus (n=3), a virus reported in dogs as well as cats [27–31], might similarly reflect environmental contamination from local pets.

More frequent infections with anelloviruses were detected in the more isolated villages of Seywiaka and Umandita. Anellovirus concentration in blood are highly dependent on the host’s immunocompetence and viral titers have been shown to increase in febrile patients [32], immune-suppressed transplant patients [33–35] and AIDS patients [36–41]. The higher rate of detectable anellovirus infections in the most isolated villages may therefore reflect generally weaker immune systems leading to more readily detectable anelloviruses possibly a result of poorer diet and medical care in these remote locations. The converse relationship was found for papillomaviruses which were more commonly detected in the most exposed village of Calabazo (21 distinct infections) versus the more isolated villages (10 and 6 infections in Seywiaka and Umandita respectively). Carcinogenic papillomaviruses were not detected. The number of papillomavirus infections therefore appears to correlate with the amount of exposure to people from outside their villages and was the only virus family where geographical isolation was associated with reduced viral diversity. Whether papillomavirus infections are consistently lower in prevalence in other small, isolated, villages relative to more connected or larger populations, remains to be further tested and confirmed.

We therefore show here that children from both connected and highly isolated villages in Northern Colombia carry diverse human viruses in their nasal mucosa, most frequently anelloviruses and papillomaviruses, that rhinovirus transmission clusters can be readily detected within these small communities, and that extreme geographical and cultural isolation did not result in a general reduction in viral diversity.

Closely related picornaviruses (and caliciviruses) have also been described in fecal samples from children of highly isolated Amazonian villages in Venezuela. This observation likely also reflect ongoing transmission chains among epidemiologically linked children within very small communities as described here [42]. This recent study of fecal viromes also showed that extreme isolation did not reduce the diversity of circulating enteric viruses [42] as we show here for the nasal mucosa. These results support our conclusion that the current reach of common human viruses extends to some of the most geographically remote populations.

## Material Method

### Study population and Study design

Nasal swab samples were collected from September 2016 to February 2017 from children with no apparent clinical signs enrolled in an influenza surveillance study located in three different villages in the Magdalena Department of Colombia by the Caribbean sea (Figure 1). Nasal swabs from 21 children from each village were collected totaling 63 samples from 34 girls and 29 boys (Table 1). Dry sterile swabs (Nylon flocked, Fisher) were used in both nostril and stored in 1 ml universal transport medium (Quidel). Samples where kept on ice for 4-7 days and then stored at −80C.

### Viral metagenomics

To reduce possible batch effects, samples from the 3 locations were processed in an interdigitated manner (First sample from village 1,2,3, then second sample from village 1,2,3, then repeat) using two Illumina MiSeq runs. Individual swab supernatants (150ul) were filtered using a 0.45-µm filter (Millipore). The filtrates were treated with a mixture of DNases (Turbo DNase [Ambion], Baseline-ZERO [Epicentre], benzonase [Novagen]) and RNase (Fermentas) at 37°C for 90 minutes to enrich for viral capsid-protected nucleic acids were then extracted using a Maxwell 16 automated extractor (Promega)[43]. Random RT-PCR followed by Nextera™ XT Sample Preparation Kit (Illumina) were used to generate a library for Illumina MiSeq (2 × 250 bases) with dual barcoding as previously described[44].

### Bioinformatic analysis

An in-house analysis pipeline was used to analyze sequence data. Before analyzing raw data was pre-processed by subtracting human and bacterial sequences, duplicate sequences, and low quality reads. Following de novo assembly using the Ensemble program [45], both contigs and singlets viral sequences were then analyzed using translated protein sequence similarity search (BLASTx v.2.2.7) to all annotated viral proteins available in GenBank. Candidate viral hits were then compared to a non-virus non-redundant (nr) protein database to remove false positive viral hits. To align reads and contigs to reference viral genomes from GenBank and to generate complete or partial genome sequences the Geneious R10 program was used. For plotting read numbers to different viruses the number of reads with BLASTx E score <10^−10^ to named viruses was divided by the total number of reads multiplied by 10^4^ then log 10 transformed to determine the size of the colored circles using Excel.

### Phylogenetic analyses

Phylogenetic trees were constructed from VP4-VP2 nucleotide sequence for rhinoviruses and amino acid sequence for papillomaviruses. Evolutionary analyses were conducted in MEGA6 using the using the Maximum Likelihood method based on the General Time Reversible model [46,47].

### Statistical methods

To evaluate the proportional distribution of viral types among villages, a nonparametric, oneway, ANOVA was performed using the Kruskal Wallis test with ties and an a priori statistical significance level set at p<0.05. Stata/MP 15.1 (StataCorp, College Station, TX) was used for the statistical analysis. The Kruskal-Wallis equality of population rank test was done using two degree of freedom.

### Ethics statement

Studies were approved by the Indigenous Health Council, Tropical Health Foundation Ethics Committee, and St. Jude Children’s Research Hospital Institutional Review Board. The investigators ensure that this study is conducted in full conformity with the principles set forth in The Belmont Report: Ethical Principles and Guidelines for the Protection of Human Subjects of Research of the US National Commission for the Protection of Human Subjects of Biomedical and Behavioral Research (April 18, 1979) and codified in 45 CFR Part 46, 21 CFR 50, 21 CFR 56 and/or the ICH E6; 62 Federal Regulations 25691 (1997), if applicable. The investigator’s Institution’s hold current Federal Wide Assurances (FWA) issued by the Office of Human Research Protection (OHRP) for federally funded research.

## Acknowledgments

Funding sources consisted of support from Vitalant Inc. to ED and ALSAC and NIAID contract HHSN272201400006C to SSC.

## References

1. Black FL (1975) Infectious diseases in primitive societies. Science 187 (4176):515–518

2. O’Fallon BD, Fehren-Schmitz L (2011) Native Americans experienced a strong population bottleneck coincident with European contact. Proc Natl Acad Sci U S A 108 (51):20444–20448. doi:10.1073/pnas.1112563108

3. Walker RS, Sattenspiel L, Hill KR (2015) Mortality from contact-related epidemics among indigenous populations in Greater Amazonia. Sci Rep 5:14032. doi:10.1038/srep14032

4. Feng H, Shuda M, Chang Y, Moore PS (2008) Clonal integration of a polyomavirus in human Merkel cell carcinoma. Science 319 (5866):1096–1100. doi:10.1126/science.1152586

5. Siebrasse EA, Reyes A, Lim ES, Zhao G, Mkakosya RS, Manary MJ, Gordon JI, Wang D (2012) Identification of MW polyomavirus, a novel polyomavirus in human stool. J Virol 86 (19):10321–10326. doi:10.1128/JVI.01210-12

6. Yu G, Greninger AL, Isa P, Phan TG, Martinez MA, de la Luz Sanchez M, Contreras JF, Santos-Preciado JI, Parsonnet J, Miller S, DeRisi JL, Delwart E, Arias CF, Chiu CY (2012) Discovery of a novel polyomavirus in acute diarrheal samples from children. PLoS ONE 7 (11):e49449. doi:10.1371/journal.pone.0049449

7. Lim ES, Reyes A, Antonio M, Saha D, Ikumapayi UN, Adeyemi M, Stine OC, Skelton R, Brennan DC, Mkakosya RS, Manary MJ, Gordon JI, Wang D (2013) Discovery of STL polyomavirus, a polyomavirus of ancestral recombinant origin that encodes a unique T antigen by alternative splicing. Virology 436 (2):295–303. doi:10.1016/j.virol.2012.12.005

8. Yang J, Yang F, Ren L, Xiong Z, Wu Z, Dong J, Sun L, Zhang T, Hu Y, Du J, Wang J, Jin Q (2011) Unbiased parallel detection of viral pathogens in clinical samples by use of a metagenomic approach. J Clin Microbiol 49 (10):3463–3469. doi:10.1128/JCM.00273-11

9. Lysholm F, Wetterbom A, Lindau C, Darban H, Bjerkner A, Fahlander K, Lindberg AM, Persson B, Allander T, Andersson B (2012) Characterization of the viral microbiome in patients with severe lower respiratory tract infections, using metagenomic sequencing. PLoS ONE 7 (2):e30875. doi:10.1371/journal.pone.0030875

10. Bal A, Pichon M, Picard C, Casalegno JS, Valette M, Schuffenecker I, Billard L, Vallet S, Vilchez G, Cheynet V, Oriol G, Trouillet-Assant S, Gillet Y, Lina B, Brengel-Pesce K, Morfin F, Josset L (2018) Quality control implementation for universal characterization of DNA and RNA viruses in clinical respiratory samples using single metagenomic next-generation sequencing workflow. BMC Infect Dis 18 (1):537. doi:10.1186/s12879-018-3446-5

11. Graf EH, Simmon KE, Tardif KD, Hymas W, Flygare S, Eilbeck K, Yandell M, Schlaberg R (2016) Unbiased Detection of Respiratory Viruses by Use of RNA Sequencing-Based Metagenomics: a Systematic Comparison to a Commercial PCR Panel. J Clin Microbiol 54 (4):1000–1007. doi:10.1128/JCM.03060-15

12. Schlaberg R, Queen K, Simmon K, Tardif K, Stockmann C, Flygare S, Kennedy B, Voelkerding K, Bramley A, Zhang J, Eilbeck K, Yandell M, Jain S, Pavia AT, Tong S, Ampofo K (2017) Viral Pathogen Detection by Metagenomics and Pan-Viral Group Polymerase Chain Reaction in Children With Pneumonia Lacking Identifiable Etiology. J Infect Dis 215 (9):1407–1415. doi:10.1093/infdis/jix148

13. Xu L, Zhu Y, Ren L, Xu B, Liu C, Xie Z, Shen K (2017) Characterization of the Nasopharyngeal Viral Microbiome from Children with Community-Acquired Pneumonia but Negative for Luminex xTAG Respiratory Viral Panel Assay Detection. J Med Virol. doi:10.1002/jmv.24895

14. Lewandowska DW, Schreiber PW, Schuurmans MM, Ruehe B, Zagordi O, Bayard C, Greiner M, Geissberger FD, Capaul R, Zbinden A, Boni J, Benden C, Mueller NJ, Trkola A, Huber M (2017) Metagenomic sequencing complements routine diagnostics in identifying viral pathogens in lung transplant recipients with unknown etiology of respiratory infection. PLoS ONE 12 (5):e0177340. doi:10.1371/journal.pone.0177340

15. Zinter MS, Dvorak CC, Mayday MY, Iwanaga K, Ly NP, McGarry ME, Church GD, Faricy LE, Rowan CM, Hume JR, Steiner ME, Crawford ED, Langelier C, Kalantar K, Chow ED, Miller S, Shimano K, Melton A, Yanik GA, Sapru A, DeRisi JL (2018) Pulmonary Metagenomic Sequencing Suggests Missed Infections in Immunocompromised Children. Clin Infect Dis. doi:10.1093/cid/ciy802

16. Blauwkamp TA, Thair S, Rosen MJ, Blair L, Lindner MS, Vilfan ID, Kawli T, Christians FC, Venkatasubrahmanyam S, Wall GD, Cheung A, Rogers ZN, Meshulam-Simon G, Huijse L, Balakrishnan S, Quinn JV, Hollemon D, Hong DK, Vaughn ML, Kertesz M, Bercovici S, Wilber JC, Yang S (2019) Analytical and clinical validation of a microbial cell-free DNA sequencing test for infectious disease. Nature Microbiology. doi:10.1038/s41564-018-0349-6

17. Gu W, Miller S, Chiu CY (2019) Clinical Metagenomic Next-Generation Sequencing for Pathogen Detection. Annu Rev Pathol 14:319–338. doi:10.1146/annurev-pathmechdis-012418-012751

18. Parize P, Muth E, Richaud C, Gratigny M, Pilmis B, Lamamy A, Mainardi JL, Cheval J, de Visser L, Jagorel F, Ben Yahia L, Bamba G, Dubois M, Join-Lambert O, Leruez-Ville M, Nassif X, Lefort A, Lanternier F, Suarez F, Lortholary O, Lecuit M, Eloit M (2017) Untargeted next-generation sequencing-based first-line diagnosis of infection in immunocompromised adults: a multicentre, blinded, prospective study. Clin Microbiol Infect 23 (8):574 e571–574 e576. doi:10.1016/j.cmi.2017.02.006

19. Schlaberg R, Chiu CY, Miller S, Procop GW, Weinstock G, Professional Practice C, Committee on Laboratory Practices of the American Society for M, Microbiology Resource Committee of the College of American P (2017) Validation of Metagenomic Next-Generation Sequencing Tests for Universal Pathogen Detection. Arch Pathol Lab Med 141 (6):776–786. doi:10.5858/arpa.2016-0539-RA

20. Wylie KM, Mihindukulasuriya KA, Zhou Y, Sodergren E, Storch GA, Weinstock GM (2014) Metagenomic analysis of double-stranded DNA viruses in healthy adults. BMC Biol 12:71. doi:10.1186/s12915-014-0071-7

21. Goggin RK, Bennett CA, Bassiouni A, Bialasiewicz S, Vreugde S, Wormald PJ, Psaltis AJ (2018) Comparative Viral Sampling in the Sinonasal Passages; Different Viruses at Different Sites. Frontiers in cellular and infection microbiology 8:334. doi:10.3389/fcimb.2018.00334

22. Bogaert D, Keijser B, Huse S, Rossen J, Veenhoven R, van Gils E, Bruin J, Montijn R, Bonten M, Sanders E (2011) Variability and diversity of nasopharyngeal microbiota in children: a metagenomic analysis. PLoS ONE 6 (2):e17035. doi:10.1371/journal.pone.0017035

23. Fry AM, Lu X, Olsen SJ, Chittaganpitch M, Sawatwong P, Chantra S, Baggett HC, Erdman D (2011) Human rhinovirus infections in rural Thailand: epidemiological evidence for rhinovirus as both pathogen and bystander. PLoS ONE 6 (3):e17780. doi:10.1371/journal.pone.0017780

24. Iwane MK, Prill MM, Lu X, Miller EK, Edwards KM, Hall CB, Griffin MR, Staat MA, Anderson LJ, Williams JV, Weinberg GA, Ali A, Szilagyi PG, Zhu Y, Erdman DD (2011) Human rhinovirus species associated with hospitalizations for acute respiratory illness in young US children. J Infect Dis 204 (11):1702–1710. doi:10.1093/infdis/jir634

25. Singleton RJ, Bulkow LR, Miernyk K, DeByle C, Pruitt L, Hummel KB, Bruden D, Englund JA, Anderson LJ, Lucher L, Holman RC, Hennessy TW (2010) Viral respiratory infections in hospitalized and community control children in Alaska. J Med Virol 82 (7):1282–1290. doi:10.1002/jmv.21790

26. van Benten I, Koopman L, Niesters B, Hop W, van Middelkoop B, de Waal L, van Drunen K, Osterhaus A, Neijens H, Fokkens W (2003) Predominance of rhinovirus in the nose of symptomatic and asymptomatic infants. Pediatric allergy and immunology: official publication of the European Society of Pediatric Allergy and Immunology 14 (5):363–370

27. Kapoor A, Mehta N, Dubovi EJ, Simmonds P, Govindasamy L, Medina JL, Street C, Shields S, Lipkin WI (2012) Characterization of novel canine bocaviruses and their association with respiratory disease. J Gen Virol 93 (Pt 2):341–346. doi:10.1099/vir.0.036624-0

28. Piewbang C, Jo WK, Puff C, Ludlow M, van der Vries E, Banlunara W, Rungsipipat A, Kruppa J, Jung K, Techangamsuwan S, Baumgartner W, Osterhaus A (2018) Canine Bocavirus Type 2 Infection Associated With Intestinal Lesions. Vet Pathol 55 (3):434–441. doi:10.1177/0300985818755253

29. Bodewes R, Lapp S, Hahn K, Habierski A, Forster C, Konig M, Wohlsein P, Osterhaus AD, Baumgartner W (2014) Novel canine bocavirus strain associated with severe enteritis in a dog litter. Vet Microbiol 174 (1-2):1–8. doi:10.1016/j.vetmic.2014.08.025

30. Niu J, Yi S, Wang H, Dong G, Zhao Y, Guo Y, Dong H, Wang K, Hu G (2019) Complete genome sequence analysis of canine bocavirus 1 identified for the first time in domestic cats. Arch Virol 164 (2):601–605. doi:10.1007/s00705-018-4096-z

31. Lau SK, Woo PC, Yeung HC, Teng JL, Wu Y, Bai R, Fan RY, Chan KH, Yuen KY (2012) Identification and characterization of bocaviruses in cats and dogs reveals a novel feline bocavirus and a novel genetic group of canine bocavirus. J Gen Virol 93 (Pt 7):1573–1582. doi:10.1099/vir.0.042531-0

32. McElvania TeKippe E, Wylie KM, Deych E, Sodergren E, Weinstock G, Storch GA (2012) Increased prevalence of anellovirus in pediatric patients with fever. PLoS ONE 7 (11):e50937. doi:10.1371/journal.pone.0050937

33. Young JC, Chehoud C, Bittinger K, Bailey A, Diamond JM, Cantu E, Haas AR, Abbas A, Frye L, Christie JD, Bushman FD, Collman RG (2015) Viral metagenomics reveal blooms of anelloviruses in the respiratory tract of lung transplant recipients. American journal of transplantation: official journal of the American Society of Transplantation and the American Society of Transplant Surgeons 15 (1):200–209. doi:10.1111/ajt.13031

34. De Vlaminck I, Khush KK, Strehl C, Kohli B, Luikart H, Neff NF, Okamoto J, Snyder TM, Cornfield DN, Nicolls MR, Weill D, Bernstein D, Valantine HA, Quake SR (2013) Temporal response of the human virome to immunosuppression and antiviral therapy. Cell 155 (5):1178–1187. doi:10.1016/j.cell.2013.10.034

35. Blatter JA, Sweet SC, Conrad C, Danziger-Isakov LA, Faro A, Goldfarb SB, Hayes D, Jr., Melicoff E, Schecter M, Storch G, Visner GA, Williams NM, Wang D (2018) Anellovirus loads are associated with outcomes in pediatric lung transplantation. Pediatr Transplant 22 (1). doi:10.1111/petr.13069

36. Li L, Deng X, Linsuwanon P, Bangsberg D, Bwana MB, Hunt P, Martin JN, Deeks SG, Delwart E (2013) AIDS alters the commensal plasma virome. J Virol 87 (19):10912–10915. doi:10.1128/JVI.01839-13

37. Sherman KE, Rouster SD, Feinberg J (2001) Prevalence and genotypic variability of TTV in HIV-infected patients. Digestive diseases and sciences 46 (11):2401–2407

38. Touinssi M, Gallian P, Biagini P, Attoui H, Vialettes B, Berland Y, Tamalet C, Dhiver C, Ravaux I, De Micco P, De Lamballerie X (2001) TT virus infection: prevalence of elevated viraemia and arguments for the immune control of viral load. Journal of clinical virology: the official publication of the Pan American Society for Clinical Virology 21 (2):135–141

39. Shibayama T, Masuda G, Ajisawa A, Takahashi M, Nishizawa T, Tsuda F, Okamoto H (2001) Inverse relationship between the titre of TT virus DNA and the CD4 cell count in patients infected with HIV. Aids 15 (5):563–570

40. Thom K, Petrik J (2007) Progression towards AIDS leads to increased Torque teno virus and Torque teno minivirus titers in tissues of HIV infected individuals. J Med Virol 79 (1):1–7. doi:10.1002/jmv.20756

41. Christensen JK, Eugen-Olsen J, M SL, Ullum H, Gjedde SB, Pedersen BK, Nielsen JO, Krogsgaard K (2000) Prevalence and prognostic significance of infection with TT virus in patients infected with human immunodeficiency virus. J Infect Dis 181 (5):1796–1799. doi:10.1086/315440

42. Siqueira JD, Dominguez-Bello MG, Contreras M, Lander O, Caballero-Arias H, Xutao D, Noya-Alarcon O, Delwart E (2018) Complex virome in feces from Amerindian children in isolated Amazonian villages. Nat Commun 9 (1):4270. doi:10.1038/s41467-018-06502-9

43. Victoria JG, Kapoor A, Li L, Blinkova O, Slikas B, Wang C, Naeem A, Zaidi S, Delwart E (2009) Metagenomic analyses of viruses in stool samples from children with acute flaccid paralysis. J Virol 83 (9):4642–4651. doi:10.1128/JVI.02301-08

44. Li L, Deng X, Mee ET, Collot-Teixeira S, Anderson R, Schepelmann S, Minor PD, Delwart E (2015) Comparing viral metagenomics methods using a highly multiplexed human viral pathogens reagent. J Virol Methods 213:139–146. doi:10.1016/j.jviromet.2014.12.002

45. Deng X, Naccache SN, Ng T, Federman S, Li L, Chiu CY, Delwart EL (2015) An ensemble strategy that significantly improves de novo assembly of microbial genomes from metagenomic next-generation sequencing data. Nucleic Acids Res. doi:10.1093/nar/gkv002

46. S. NMaK (2000) Molecular Evolution and Phylogenetics. Oxford University Press, New York.

47. Tamura K, Stecher G, Peterson D, Filipski A, Kumar S (2013) MEGA6: Molecular Evolutionary Genetics Analysis version 6.0. Mol Biol Evol 30 (12):2725–2729. doi:10.1093/molbev/mst197

